# Manual versus automatic annotation of transposable elements: case studies in *Drosophila melanogaster* and *Aedes albopictus*, balancing accuracy and biological relevance

**DOI:** 10.1101/2025.01.10.632341

**Authors:** Tomas Carrasco-Valenzuela, Alba Marino, Jessica M. Storer, Iago Bonnici, Camila J. Mazzoni, Michael C. Fontaine, Annabelle Haudry, Matthieu Boulesteix, Anna-Sophie Fiston-Lavier

## Abstract

Transposable elements (TEs) play a pivotal role in genome evolution, yet their detection and annotation remain challenging due to the limitations of current methods. Manual curation is considered the gold standard for generating TE libraries, particularly for TE focused studies, although it requires extensive training and time. With the rapid increase in genome assembly publications and the growing need for large-scale comparative analyses, automated software for TE annotation has become indispensable. This study compares manual and automated approaches to TE detection and annotation, focusing on two species: *Drosophila melanogaster* and *Aedes albopictus*.

In *D. melanogaster*, a species with a well-annotated TE repertoire and a smaller genome, the differences between manual curation (MCTE) and automated annotation (ATTE) are relatively minor. However, significant differences arise when analysing *Ae. albopictus*, a species with a larger genome and higher TE diversity. While automated methods identified a greater number of TEs, including many smaller and fragmented elements, manual curation provided more detailed classifications and on average larger consensi. Automated pipelines offer a viable alternative for genome-wide analyses such as TE content estimate, particularly when time and resources are limited. However, caution is advised when interpreting results, as finer details of TE dynamics may be overlooked.

This study highlights that the choice of annotation method depends on the intended analysis. Manual curation is more suitable for TE population genomics and studies focusing on recent transposable element activity, while automated methods are appropriate for larger comparative analyses or genome assembly projects. Ultimately, both methods have their strengths and limitations, and understanding the specific features of the genome and repeatome under study is essential for selecting the appropriate approach.

## Introduction

As high-quality genome assemblies become more widely available, the analysis and annotation of these genomes present a significant challenge. This is particularly true for identifying transposable elements (TEs), which are highly mutagenic and prone to losing their sequence identity over time and across different genomes (McDonald, Weinstein, and Lambert 1988; Arkhipova 2018). TEs are major contributors to genome evolution, gene organisation and expression; thus, their proper identification is crucial for understanding their roles in adaptation, as well as in the conflicts and cooperation with their host genomes (Hayward and Gilbert 2022; Casacuberta and González 2013).

Several pipelines, such as RepeatModeler2 (Flynn et al. 2020), REPET (Rafii and Pardo 2013) or EDTA (Ou et al. 2019) to name a few, have been developed and are widely used by the research community to identify TEs. Despite their differences, these methods all follow a similar core principle: they search for repeated loci within the genome. Some approaches compare these loci to curated TE sequence databases (S. Orozco-Arias et al. 2023), while others generate *de novo* consensus sequences. For a deeper comparison of their differences and similarities, see Hoen et al. (2015). Ultimately, the confidence in the resulting TE library depends on the quality of the reference TE databases and the accuracy in detecting biologically meaningful repetitions in the genome.

However, automatic detection algorithms usually lack the precision and exactitude required for certain downstream applications (Goubert et al. 2022; Lerat 2010; Flynn et al. 2020). The best way to reach such quality is to curate manually each generated TE consensus: this implies checking for specific TE indicators and hallmarks, such as structural repetitive motifs and coding domains. This process is very tedious, time-consuming, and difficult to reproduce, as well as challenging for non TE-specialists (Goubert et al. 2022).

However, manual curation serves as a means of validation for computational predictions, allowing researchers to confirm the accuracy of TE annotations and refine results based on specific biological contexts. Platt et al (2016) started to define a set of recommendations for manual curation adopted by (Peona et al. 2021; Jebb et al. 2020; Louha et al. 2020) and reviewed by (J. M. Storer et al. 2021; Goubert et al. 2022).

Very recently, some tools have been developed to automate the curation process (Baril, Imrie, and Hayward 2022; S. Orozco-Arias et al. 2023; Simon Orozco-Arias et al. 2022): they are promising as they can improve the quality of automatically generated TE libraries, as well as aid manual curators in their task and prevent reproducibility issues.

Despite the evident need for manual curation when focusing on TE biology, researchers will tend to choose an automated approach over a manual one or the other way around, depending on time availability, expertise, and scope of their project. However, this choice might ultimately affect the results of downstream analyses.

Studying the contribution of TEs to recent adaptation can lead to focus on recently active and abundant TE families. Furthermore, because genotyping methods involve read mapping at the extremities of TE insertions, the description of full-length consensi is a priority for this kind of study.

Such an approach is impractical when a global assessment of the entire genome is required, and/or numerous taxa are to be annotated. On the other hand, a “simple” *de novo* annotation can lack the level of definition needed, due to redundancy, fragmentation and unclassified elements (Baril, Imrie, and Hayward 2022; Rodriguez and Makałowski 2022). A detection workflow that is time-effective and that will produce accurate TE annotations is therefore a preferable trade-off to allow in-depth comparative analysis of TE accumulation across numerous non-model species. To meet such needs, some authors of this paper chose two different approaches and independently developed two different protocols - a manual-based one and a fully automated one - to annotate the genome of the same species, the Asian tiger mosquito (*Aedes albopictus*). The former approach involves *de novo* (RepeatModeler2, EDTA, MITE-Tracker) and homology-based (RepeatMasker used with RepBase) TE discovery, followed by redundancy removal, consensi extension and extensive manual curation: its product is hereafter referred to as Manually Curated TE (MCTE) library. The latter combines the output of two identification methods (RepeatModeler2 and EDTA), to then reduce redundancy and perform automatic curation of TE sequences using MCHelper in fully automated mode: its product is hereafter referred to as Automatic Tools TE (ATTE) library.

*Aedes albopictus* genome is large by Dipteran standards (1.190–1.275 Gb; (Palatini et al. 2020); earlier studies based on sequencing data estimated that repetitive elements make up about 50%, with at least 33.58% being TE-derived (Goubert et al. 2015). More recent research based on transcriptomic data suggests that TEs comprise over 40% of the genome (de Melo and Wallau 2020). This makes it difficult to determine in advance the completeness and accuracy of both the ATTE and MCTE approaches. Therefore, as a control, we included *Drosophila melanogaster* in the analysis, given its compact genome (180 Mb), excellent genome assembly and comprehensive TE annotation (Rech et al., 2022).

Initial comparisons between the MCTE and ATTE approaches for both genomes revealed minimal overlap between the obtained libraries. We explored these discrepancies in *Ae. albopictus* and examined whether they arose from the genome’s complexity or the non-exhaustiveness of the MCTE approach. ATTE tends to identify more sequences, particularly smaller and fragmented elements, while MCTE provides longer consensi with more detailed classifications. This difference is especially evident in *Ae. albopictus,* which harbours greater TE diversity.

In comparing TE libraries across two genomes—*D. melanogaster*, a model organism with only 20% TE content, and *Ae. albopictus*, with its large genome (1.3 Gb) and ∼40% TE content—we found that both approaches produce quantitatively distinct genome annotations. While automated curation cannot fully replace manual efforts, the two methods are complementary. Automated approaches can guide manual curation or be integrated to generate more exhaustive libraries.

Finally, we provide here a composite TE library for *Ae. albopictus* by combining both methods, which can be used for future studies.

## Material and methods

### TEs discovery for manual curation in *Ae. albopictus*

For the construction of the MCTE library, TE consensi were obtained from the AalbF2 genome assembly (Aalbo_primary.1; (Palatini et al. 2020)) using three distinct annotation algorithms: EDTA-2.0.0 (Ou et al. 2019), RepeatModeler-2.0.2a (Flynn et al. 2020), and MITE-Tracker (Crescente et al. 2018). The resulting TE consensi were combined into a single multifasta file, along with a subset of sequences from RepBase 25.08 (Jurka et al. 2005; Bao, Kojima, and Kohany 2015) (including sequences from *Anopheles gambiae*, *Drosophila melanogaster*, Invrep -Invertebrata-, and Invsub -Invertebrata subfamilies-). This file was then used to mask the genome with the RepeatMasker command:

`RepeatMasker -pa 64 -s -a -inv -nolow -gff -dir. -lib $WORK_DIR/TE_Aealb.raw.fa -cutoff 250 $ASSEMBLY`. RepeatMasker hits shorter than 80 bps were removed due to the likelihood of spurious results. After masking the genome, the OneCodeToFindThemAll pipeline (Bailly-Bechet, Haudry, and Lerat 2014) was applied to reconstruct TE copies by merging close-by RepeatMasker hits using the following command:

`build_dictionary.pl --rm Aalb.out --unknown --fuzzy > dico_fuzzy.txt \ one_code_to_find_them_all.pl --rm Aalb.out --ltr dico_fuzzy.txt --fasta --flanking 100 --strict --unknown --insert 80`

A custom script was then used to parse the output of one code to find them all (Bailly-Bechet, Haudry, and Lerat 2014) and extract a fasta file containing all the TE copies sequences (https://github.com/Tcvalenzuela/Manual-versus-automatic-annotation-of-transposable-elements/octfta_to_fasta.sh).

In order to group TE copies by families, RepeatMasker-identified insertions were clustered using cd-hit (Huang et al. 2010; Li and Godzik 2006) with parameters set to match as much as possible the 80-80-80 rule : identity of 80% or greater along more than 80% of the sequence in sequences longer than 80 bp as described in Flutre (2011) - as follows: `cd-hit est -i copies.fasta -o consensi.fasta -c 0.8 -G 0 -aS 0.8 -M 90000 -d 0`.

After several test runs, in order to increase the high copy number families, we selected only clusters with more than seven insertions. Consensi were called from the remaining clusters with the Refiner tool from RepeatModeler2 (Flynn et al. 2020), after downsampling 500 sequences from the largest clusters. In order to reduce redundancy among family consensi, these were in turn clustered with cd-hit-est and consensi called again with Refiner (Flynn et al. 2020). The process was repeated until no redundancy was detected in the library (total number of iterations: 8). In the end, 23,009 family consensi were obtained at this step.

In order to increase the likelihood of obtaining full-length TE models, all 23,009 consensi were extended using the following method. First, the putative TEs were screened for low complexity and simple repeat sequences using TRF (Benson 1999). To gather sequences for extension, the filtered sequences were used as a library for RepeatMasker (v4.1.5) on the AalbF2 genome assembly. An alignment per family is generated in stockholm (.stk) format, followed by extension, using somewhat relaxed parameters for extension into the flanking sequence based on the chromosomal locations present in the .stk files. Following extension, additional identifying information might have been obtained, such as long terminal repeat (to allow for endogenous retroviruses identification), terminal inverted repeat (for DNA family identification, or polyA tail (LINE/SINE identification). Therefore, RepeatClassifier (a utility part of RepeatModeler) is run to take this additional information into account.

The extended consensi was used as a reference in a second RepeatMasker run, this time performed on AalbF3 -a deduplicated version of F2 plus optical mapping-reference genome assembly GCA_018104305.1 (Boyle et al. 2021). The output of RepeatMasker was used to sort the list of consensi by full-length insertion total number, where a full-length insertion is defined as being at least 90 percent the length of the consensus, with a maximum nucleotide divergence of 20 percent (Flutre et al. 2011). Using the custom script https://github.com/Tcvalenzuela/Manual-versus-automatic-annotation-of-transposable-elements/detectFullSize.py, the list of consensi was finally sorted by full-length frequency and manual curation was done starting from the most frequent consensi, elements with less than three insertions were not considered for manual curation and were not part of the final library neither.

### Manual curation of TEs from *Ae. albopictus*

Relevant information necessary for manual curations such as the absolute frequency of insertions, pre- and post-extension length, insertions frequency on each genome assembly (AalbF3 and AalbF5 versions), ratio of extended consensus length over original consensus length, extended consensus coverage, extended consensus median coverage, insertion frequency, full-length insertion frequency, among others (Supplementary Table 1) were computed for all the consensi. The frequency of full-length insertion was used as a priority list for the manual curation.

Manual curation was conducted on the TE consensus of the 800 most frequent full-length insertions, resulting in a final TE library containing 497 TEs. First, from the insertions coordinates from RepeatMasker and using the custom script https://github.com/Tcvalenzuela/Manual-versus-automatic-annotation-of-transposable-elements/GetMultipleAln.sh, each insertion was extended to 2,000 bp on both flanks, extracted and the 100 longest insertions clustered together using ClustalO (Sievers and Higgins 2014) for examination of the characteristic component of the respective category of TEs. Additionally, the examination of each TE consensus was done using a set of manual curation identification tools as TE-Aid (Goubert et al. 2022), RepeatClassifier (Flynn et al. 2020) and alignment visualisation using Aliview (Larsson 2014) together with databases Repbase (Jurka et al. 2005; Kohany et al. 2006; Vladimir V. Kapitonov and Jurka 2008) and CDD protein domains (Lu et al. 2020; Marchler-Bauer et al. 2015), following the annotation recommendations from Goubert et al. (2022).

In summary, following the guidelines of Wicker et al (2007): long terminal repeats (LTRs) are identified based on the presence of open reading frames (ORFs) encoding proteins such as capsid protein, aspartic proteinase, integrase, RNase H, reverse transcriptase, and envelope protein, depending on the superfamily. Additionally, the LTR element must be flanked by Target Site Duplications (TSDs) of 4–6 base pairs (Neumann et al. 2019).

Penelope-like elements can be further distinguished from homing endonucleases (ENs) by the presence of a conserved CCHH Zn-finger motif, where two cysteines are located between the GIY and YIG motifs (Arkhipova 2006). For LINEs, the typical structure includes a retrotranscriptase combined with an apurinic endonuclease or ENs domain, and some families have an additional ORF containing RNase H (Wicker et al. 2007). For LINEs and Penelope-like elements, 5’ truncation products are expected on clustering of different insertions <u>(Arkhipova 2006)</u>.

Terminal Inverted Repeats (TIRs) and variable-size TSDs are key identifiers for most class II elements. Due to the absence of conserved protein domains across all class II transposons, the identification of Class II elements rely on Terminal inverted repeats (TIR) and target site duplication (TSD). After the identification of TIR and TSD, their open reading frames (ORF) phylogeny is critical for classification between Class II families. In eukaryotic transposons, the transposase encoded is a DDD/E transposase (Kojima 2020). For non-autonomous copies, it is important to check if the TIR matches known autonomous elements. If no match is found and the consensus sequence is shorter than 800 bp, the element is classified as a MITE (Hu, Zheng, and Shang 2018). If it exceeds 800 bp, it is categorised as ClassII_other.

More complex class II elements, such as Helitrons and Mavericks, can be classified based on homology to protein domains due to their unique protein sets and lack of TSDs (V. V. Kapitonov and Jurka 2001) although most of these elements are non-autonomous.

While LTRs and TIRs should be detectable using TE-Aid, there are instances where TIRs are not immediately visible, even when coverage suggests a class II element. In such cases, manual inspection using a visualizer like Aliview (Larsson 2014) to locate the TIR is recommended.

As in previous analyses on TEs on mosquitos (de Melo and Wallau 2020), there are limitations to the TE annotation due to lack of homology to TE databases or to the loss of ORF due to accumulated mutations. For some elements, it was impossible to get further than Class I or Class II classification in both MCTE and ATTE library construction. To simplify the comparison, these elements are referred to as ClassI_other and ClassII_other.

### Manually annotated library for *D. melanogaster*

The evaluation of the ATTE method also depends on the completeness of the MCTE library used as a control. Therefore, for *D. melanogaster*, we used the manually curated TE library from Rech et al. (2022). They used long-read data to assemble high-quality genomes of 32 *D. melanogaster* natural strains from five different climatic regions. From this genomic resource, Rech et al. (2022) manually curated a TE dataset composed of 165 consensus sequences, thoroughly representing the genomic TE diversity in this species.

### Automated discovery and curation of TEs in *Ae. albopictus* and *D. melanogaster*

The study of TE dynamics in larger numbers of species can require an automated strategy of TE discovery where automatic quality-check and curation steps are integrated to obtain better-quality annotations. The genomes AalbF5 for *Ae. albopictus* and dm6 for *D. melanogaster* (Hoskins et al. 2015) were annotated with the following automated pipeline to obtain the respective ATTE libraries. New insertions were identified with Repeatmodeler2 (Flynn et al. 2020) and EDTA (Ou et al. 2019). Each program was run independently, and the mined sequences were concatenated on a single multifasta file. Afterward, MCHelper (S. Orozco-Arias et al. 2023) was used in fully automated mode to eliminate redundancy, remove false-positives, elongate and check for TE classification.

### Comparison of TE libraries

To compare the MCTE and ATTE libraries, we performed a clusterization of the consensi between libraries for each species following an 80-80 rule (Flutre et al. 2011), using cd-hit as follows:

~~~
cd-hit-est-2d -i <Library MC> -i2 <Library AT> -o <Output Name> -c 0.8 -G 0 -aS 0.8 -n 5 -d 0
~~~

To further compare the MCTE and ATTE libraries, we used the script “get_family_summary_paper.sh” from (Flynn et al. 2020). Briefly, a target library is masked with a reference library using RepeatMasker with -nolow option, and its level of completeness is evaluated based on the reference. The overlap between MCTE and ATTE libraries was assessed by using the ATTE and the MCTE libraries as reference, respectively.

Based on Flynn (2020), we use an inhouse script (Link to github) where each element of the reference library was assigned to one of the following 5 categories: “Perfect”, “Good,”, “Present”, “Fragmented” and “Absent”. The first 3 categories are defined following Flynn et al. (2020). “Perfect” families are those for which one sequence in the target library matches >95% in sequence identity and coverage with a family in the reference library. “Good” families are those in which multiple overlapping target-library sequences with alignments >95% similar to the reference consensus make up >95% sequence coverage of the element from the reference library. A family is considered “Present” if one or multiple target sequences align with >80% similarity to and cover >80% of the reference consensus sequence. In addition, we consider a family to be “Fragmented” if multiple query sequences with >80% similarity cover >30% of a consensus, thus accounting for potential fragments that might match full-length families in the reference library. Otherwise, we consider a family to be “Absent”.

Finally, each library was used to annotate the respective genome and compare the genomic estimates of TE content obtained with manual and automated curation. RepeatMasker (Nishimura 2000; Smit, Hubley, and Green 2015) was run with RMblast as search-engine, and annotations were used to quantify the overall genomic TE composition and the proportion occupied by each TE consensus (Supplementary Table 2).

## Results

### TE library composition

The MCTE library for *Ae. albopictus* includes 496 consensi with up to 733 full-length insertions, with an average of 94.5 insertions in the AalbF5 genome assembly. It is composed of 135 LINEs (27.16% of the total consensi), 129 MITEs, 80 unclassified elements of Class II, 79 LTRs, 60 DNA elements, 8 *Penelope-like* elements, 4 SINEs and 1 Helitron. On the other hand, the ATTE library for *Ae. albopictus* comprises 7,782 consensi and is composed of 5,750 LTRs (73.89% of the total), 1,024 DNA elements, 169 LINEs, 795 MITEs, 1 SINE and 9 RC, as well as 33 and 1 unclassified elements of Class I and Class II, respectively (Figure 1).

**Figure 1.**
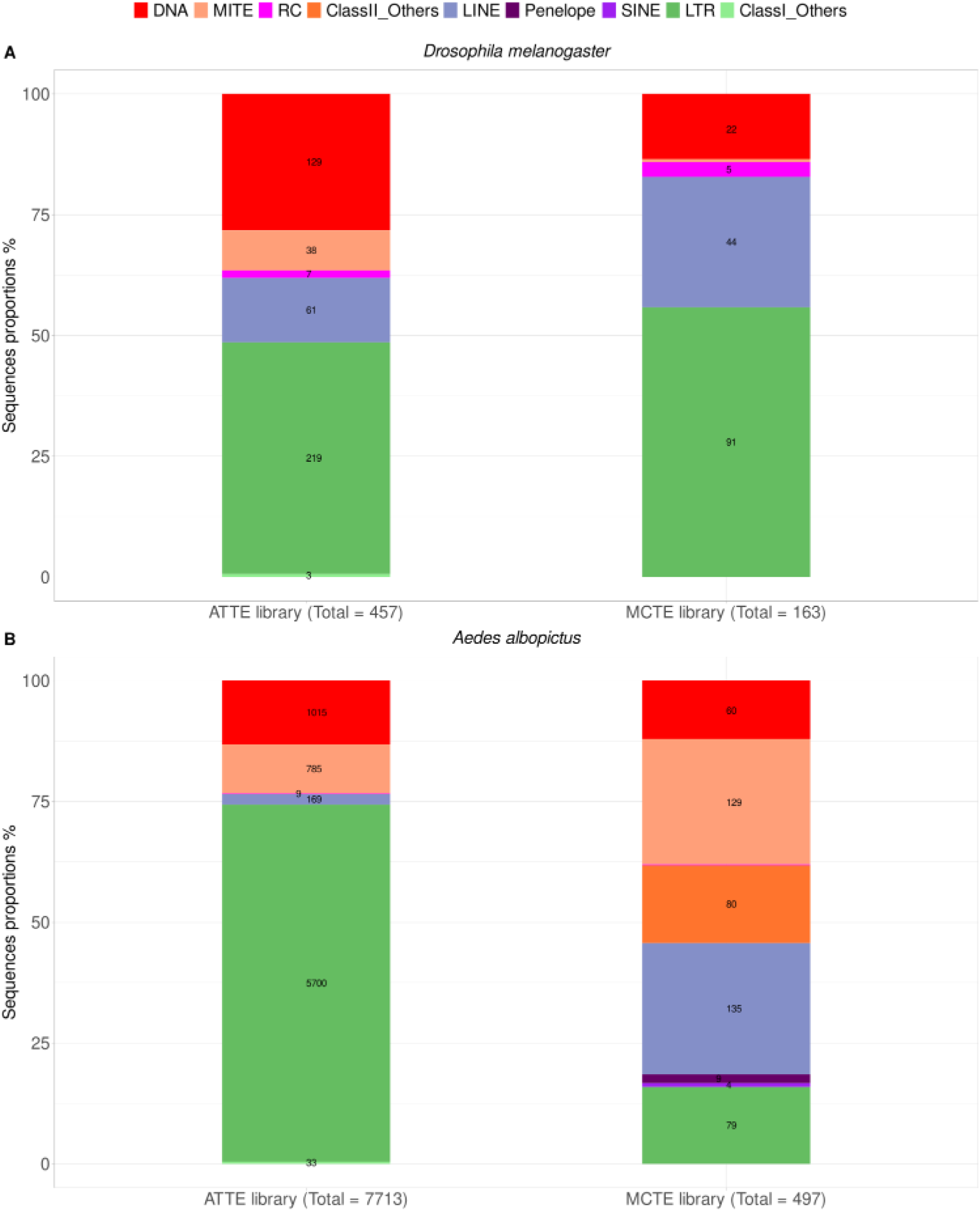
Composition of the TE libraries of (A) *D. melanogaster* and (B) *Ae. albopictus*. The proportions of TE sequences type are shown both in percentages on the y-axes and as number of sequences on the stacked bars. The number of total sequences is indicated at the bottom of barplots for each library. For categories with only one sequence, numbers are not displayed.

The MCTE library for *D. melanogaster* was obtained from the public database made available by Rech et al. (2022). Statistics including consensi length, frequency on the genome, and other descriptive statistics can be found in Supplementary Table 2. This library was composed of 165 consensi: 91 LTR elements (55.15% of the total), 45 LINEs, 23 DNA elements, five RC, and one MITE. The *D. melanogaster* ATTE library is composed of 463 consensi, and has a high proportion of LTR elements with 225 consensi (48.6% of the total consensi), 129 DNA consensi, 61 LINEs, 38 MITEs, 7 RC and 3 unclassified elements of Class I (Figure 1).

In both species, the ATTE library contains more consensi than the MCTE library, although this is particularly evident in *Ae. albopictus* where the ATTE library contains 15 times more consensi. For *D. melanogaster*, the consensus lengths of the ATTE library are not significantly shorter than those of the MCTE library for any TE order (Figure 2; Wilcoxon test: DNA: W = 1701, p = 0.138; LINE: W = 1512, p = 0.271; LTR: W = 11112, p = 0.111; MITE: W = 38, p = 0.100). In *Ae. albopictus*, LTR and LINE elements are significantly shorter in the ATTE library compared to their counterparts in the MCTE library (LTRs: average length: 2,913 bp in the ATTE library vs. 6,738 bp in the MCTE library; LINEs: average length: 3,670 bp in the ATTE library vs. 4,574 bp in the MCTE library). This difference in length is statistically significant for both LTR (Wilcoxon test: W = 66,322, p = 4.07e-27) and LINE elements (Wilcoxon test: W = 7,881, p = 3.66e-6).

**Figure 2.**
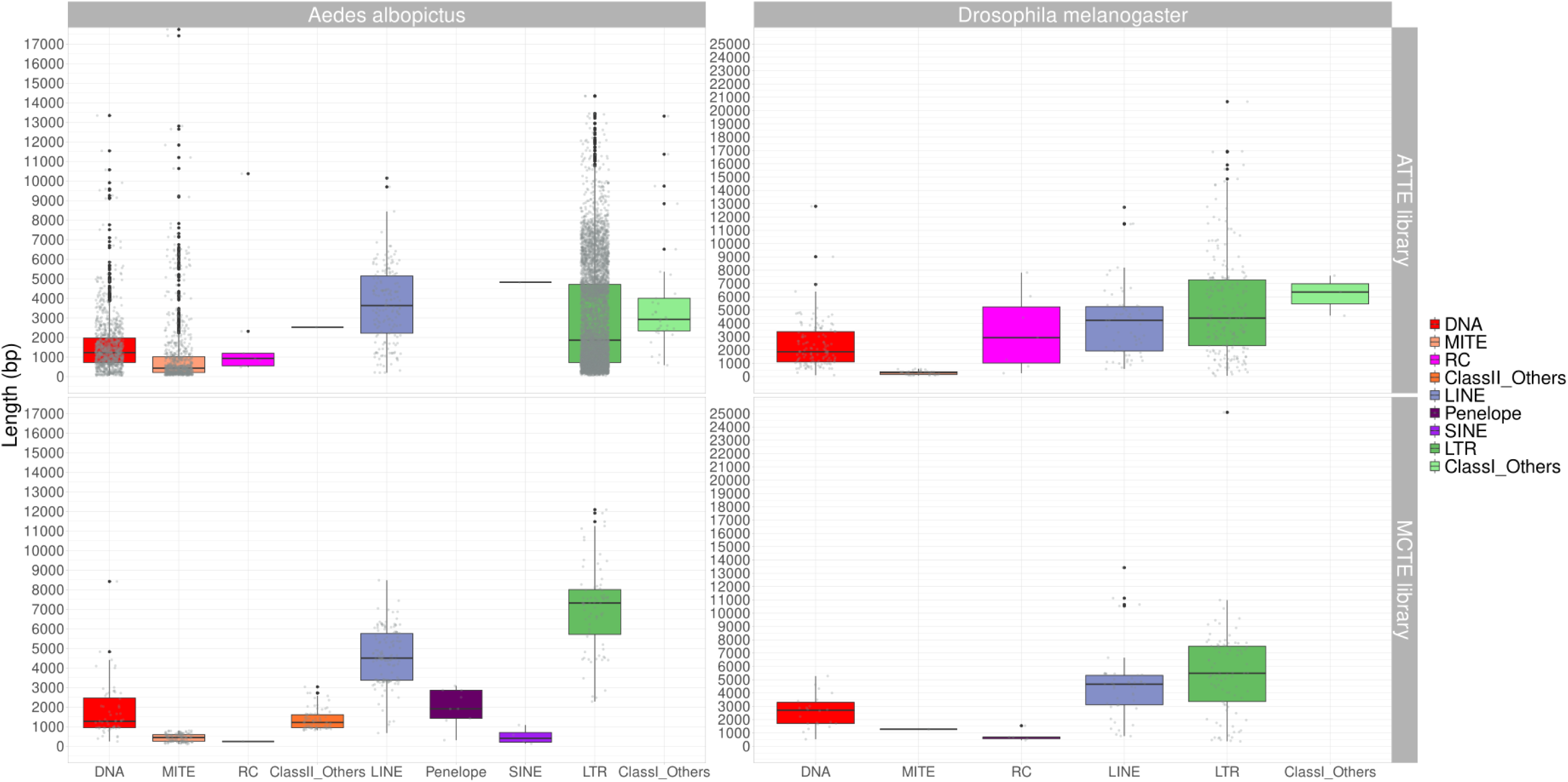
TE size distribution for MCTE (bottom panels) and ATTE libraries (top panels). For each TE library, the lengths of the consensus sequences were measured, and the distribution plotted grouping the elements by their respective TE order.

### How similar are libraries automatically generated versus those manually curated?

While having identical, complete copies from start to finish in the MCTE and ATTE libraries is ideal, achieving this for TEs is seldom possible in the current state of research. However, we can assess how overlapping MCTE and ATTE libraries are by comparing sequence similarity and coverage.

The libraries were clustered using the 80-80 rule, as detailed in the methods section. In *Ae. albopictus*, 58.63% of the consensi from the MCTE library had no corresponding match in the ATTE library (Supplementary Table 3). Of the remaining 41.37% of clustered sequences, two-thirds matched with consensi of the same TE order. In contrast, 98.10% of the ATTE library lacked a match in the MCTE library, with only 0.82% clustering with sequences of the same TE order.

Similar results were observed in D. melanogaster, where 98.27% of the ATTE library consensi (Supplementary Table 3) had no match in the MCTE library. Of the remaining 1.73% (8 consensi), 7 matched at the TE order level. On the other hand, 74.55% of the MCTE library had no equivalent consensi in the ATTE library, while the remaining 25.45% were largely composed of consensi matching the same TE order.

We compared more exhaustively the composition of the libraries by categorising consensi as “Perfect”, “Good”, “Present” (as in Flynn (2020)), and “Fragmented” and “Absent” with respect to the alternative library (see the Methods section).

Overall, 64% of the *D. melanogaster* MCTE sequences had a hit in the ATTE library and in most cases with the same TE type (Supplementary Table 4). From this, 28% of the MCTE sequences fall into the category “Perfect”, meaning that they have > 95% identity over 95% of the length with one sequence in the ATTE library (Figure 3A; Supplementary Table 4). These mostly correspond to LTR elements, followed by LINEs. “Good” sequences (those elements with >95% identity over 95% of the length in multiple sequences) make up 12% of the MCTE library and are predominantly LTRs, as well. In the reverse comparison, we identify that 75% of the sequences in the ATTE library do not have any match in the MCTE one, arguably due to the bigger size of the former (2.8 AT/MC ratio). Compared to the MCTE library, a significantly smaller fraction of the ATTE library has corresponding hits in the MCTE one (Perfect = 9%, Good = 4%, Present = 4%, Fragmented = 7%): the large majority are LTR elements and have the same classification in the two libraries.

**Figure 3.**
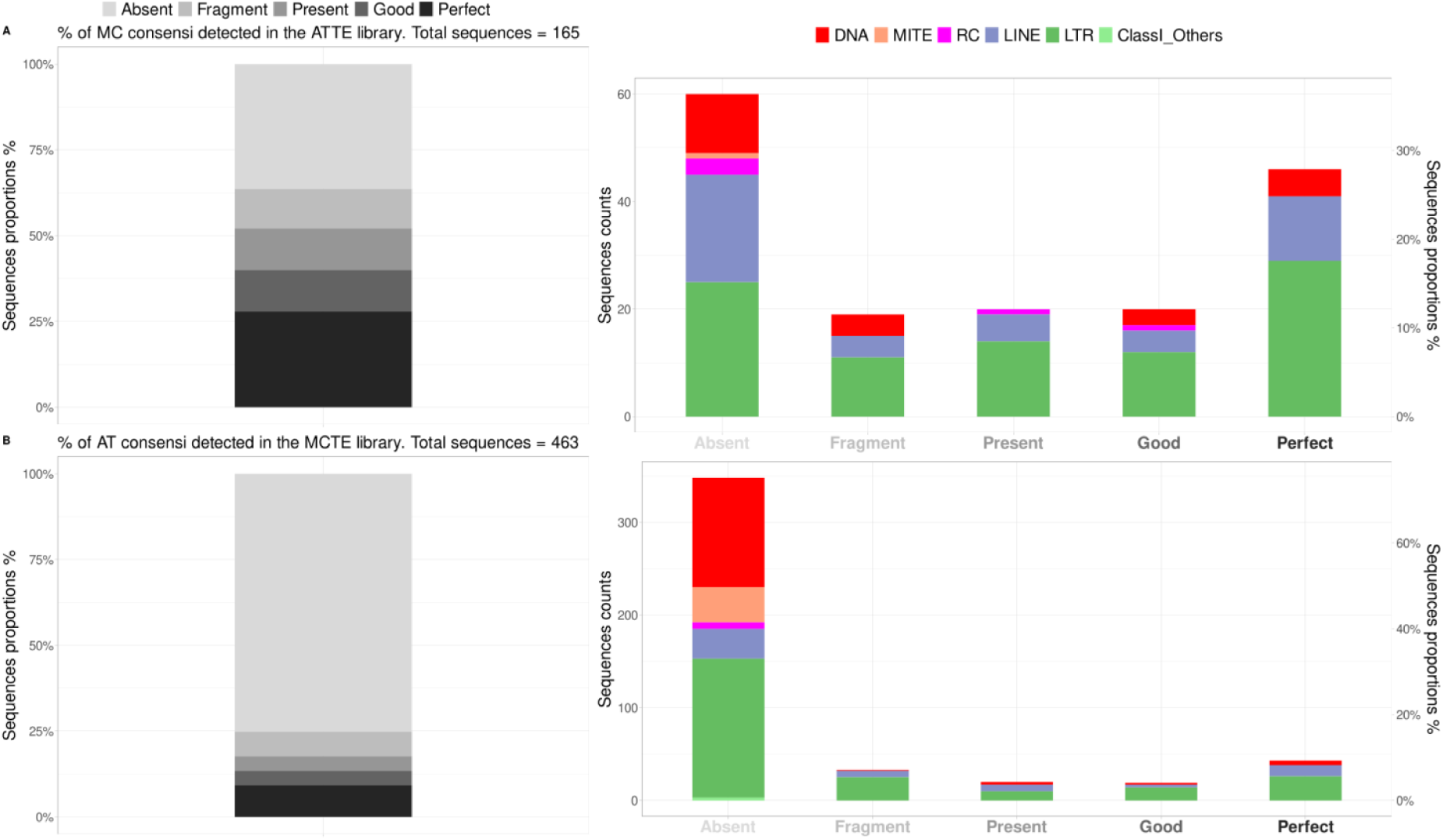
Comparison between the ATTE and MCTE libraries in *D. melanogaster*. (A) Level of completeness of the ATTE library (“query”) versus the MC library (“reference”): the percentages of TE consensi in the MCTE library detected in the ATTE library are shown for the categories Absent, Fragment, Present, Good and Perfect (left, scale of grey). For each category, the proportion of consensi is displayed by TE order (right, coloured stacked bars). (B) The same analysis was performed using the ATTE library as “reference” and the MCTE library as “query”.

The same analysis was performed on the *Ae. albopictus* libraries (Figure 4). Here, we identify that three-quarters of the MCTE elements have a similar ATTE consensus; divided on 36.82% “Perfect”, 13.88% “Good”, 16.10% “Present”, 7.65% ‘Fragmented”, and 25.55% of absent elements (Figure 4A). In particular, 59% of the MCTE consensi that have no counterpart in the AT library are LINEs. Additionally, most consensi of this category overall correspond to Class II elements including MITEs (∼25%), DNA (17%), and other non-further classified elements (23%). Furthermore, while 42.25% of the MCTE consensi matched with sequences of equivalent classification, 32.19% matched with different TE types, with MITEs constituting 46.88% of the misclassifications three times more than the next category (LINEs with 17.50% of the misclassifications) (Supplementary Table 4).

**Figure 4.**
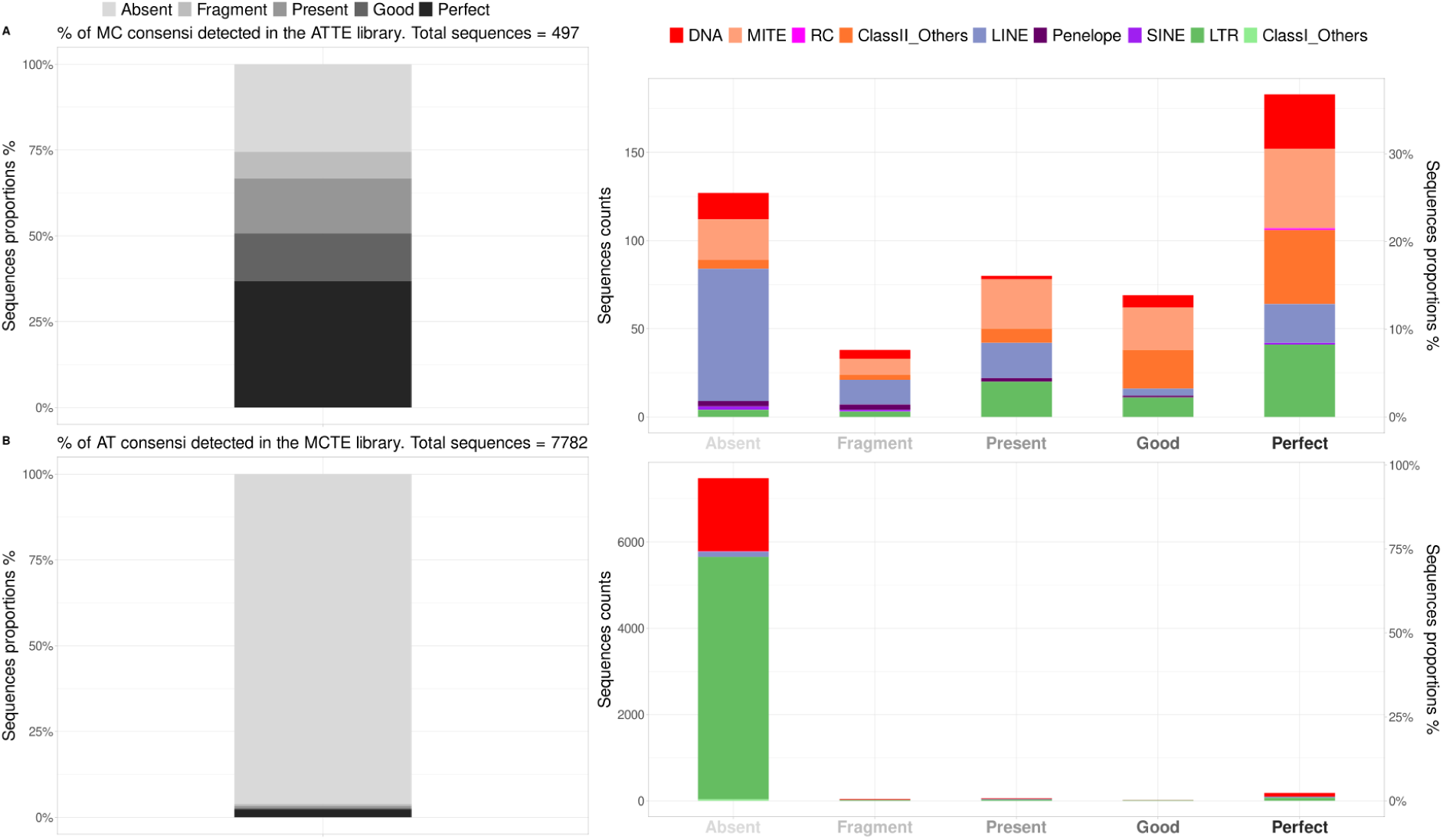
Comparison between the ATTE and MCTE libraries in *Ae. albopictus*. Level of completion for the automatically generated library versus the manually curated one in *Ae. albopictus* in panel A illustrates the percentage of TE consensi from MC (“reference”) detected in the AT (“query”) for the categories Absent, Fragment, Present, Good and Perfect in a scale of grey. For each category, the portions represented by different TE orders are displayed as stacked bars. Panel B represents the same analysis with the consensi from AT (“reference”) detected in the MC library (“query”).

On the opposite comparison (Figure 4B), we again identified a substantial number of AT consensi (96%) as absent from the MC library. This could be due to the ratio 15.5 of AT/MC consensi, hence either the overabundance of AT sequences or undersampling of MC sequences. 75% of the absent elements are LTRs, which are also the most abundant in the AT library. Out of the 302 matching AT sequences (∼ 4% of the total), 170 had hits with the same TE order; the rest had hits with sequences with different classification, notably LTR and DNA elements.

### Are we capturing the full genetic diversity of transposable elements within the genome?

The ability to draw biologically significant conclusions about TEs depends on capturing a substantial proportion of the number and diversity of TEs present in the organisms. For this, it is important to assess the genomic coverage of the different transposable detection alternatives together with the TEs variability detected. We compared the two different library strategies on the two species, masking the genomes and identifying the level of coverage per strategy as described in the methods section (Figure 5).

**Figure 5.**
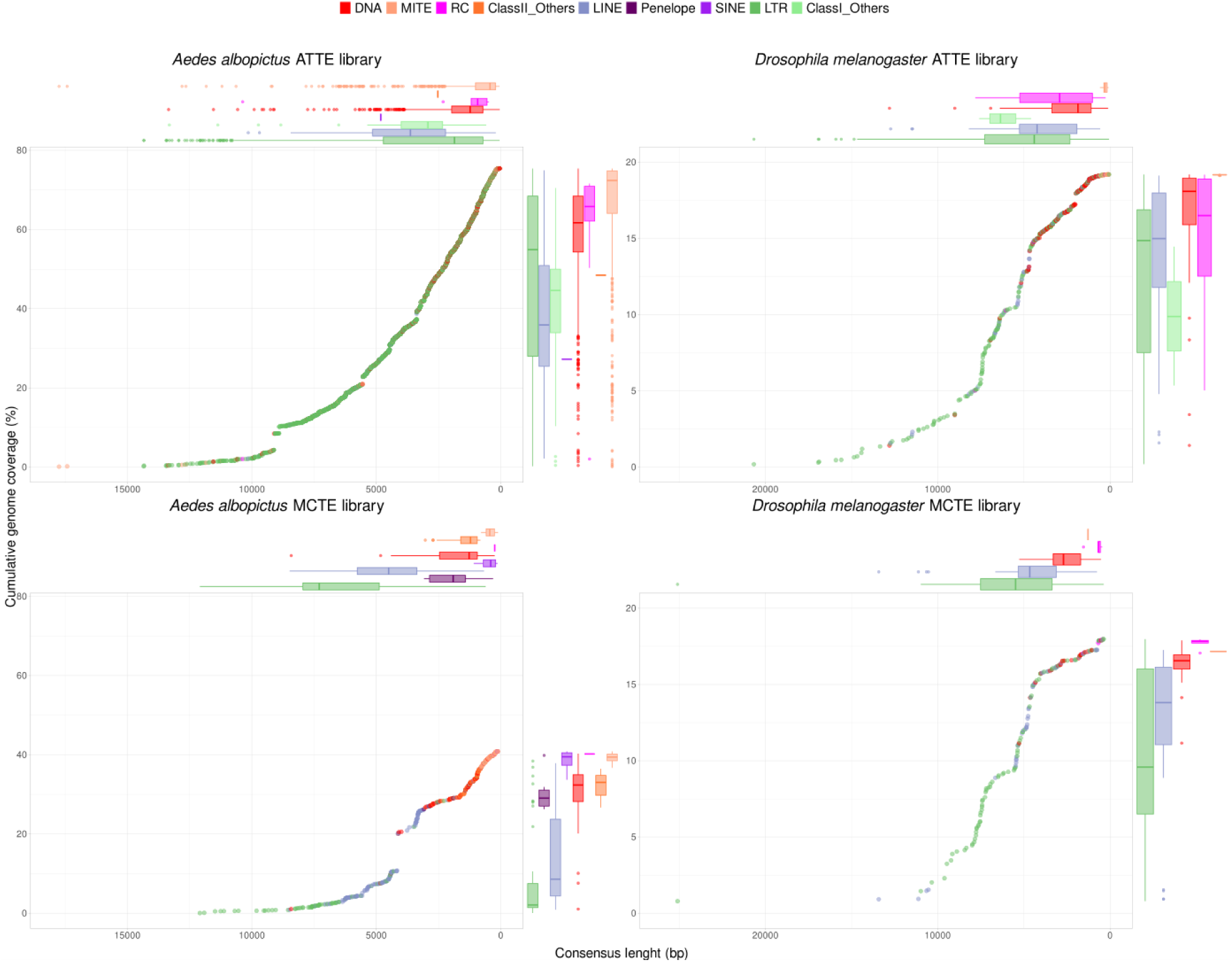
Comparative cumulative genome coverage and length distribution of TE consensi with MCTE and ATTE libraries. (A) Levels of cumulative genome coverage are shown by decreasing consensus length for *D. melanogaster* (right) and *Ae. albopictus* (left) genomes using the MCTE libraries (bottom) *versus* the ATTE libraries (top). Boxplots on the right and top of each scatterplot highlight the distribution of the different TE orders along the x- and y-axes. Colours correspond to different TE categories, as per legend on top.

For the *Ae. albopictus* genome, the TE coverage obtained with the MCTE library was lower (40.8% of the genome) than with the ATTE library (75.4% of the genome). This might be due to fragmented copies on ATTE library or a reflection of the incomplete nature of the MC library in *Ae. albopictus*, which focuses on recent TE families with >3 full-length copies in the genome. Besides this and in concomitance with results of Figure 1, we observed a prevalence of annotated LTRs with the ATTE library. Interestingly, a sharp increase in genome coverage (∼ 12 to 21%) is observed with the MCTE library: this is due to a LINE/RTE-BovB with a full-length consensus of 4,000 bp and 168,000 insertions - between fragmented and full copies - occupying 9% of the genome.

In the *Ae. albopictus* MCTE library, we observed a clear size differentiation across TE categories, reflecting the progressively smaller sizes of its elements. However, this pattern is not evident in the ATTE library for the same genome, where LTR elements dominate the entire size distribution, as shown in Figure 5.

For *D. melanogaster*, the TE genome coverage is quite similar, with 18% for MCTE and 19.2% for ATTE. This aligns with our expectations, given the non-significant differences in consensi length distribution, despite variations in sequence similarity. Moreover, the relative scarcity of TEs in *D. melanogaster* and the extensive manual curation already performed in its genome contribute to these results. Additionally, TE families in both MCTE and ATTE show less diversity and lower copy numbers compared to *Ae. albopictus*.

## Discussion

Due to the limitations of current TE detection methods, manual curation remains the gold standard for generating TE libraries, particularly for TE-focused analyses, though it requires significant training and execution time. However, with the rapid pace of genome assembly publications and the growing need for large-scale comparative analyses, automated software for TE annotation and curation is becoming increasingly indispensable. In this study, we compare manual and automated approaches, showing that differences in library composition, TE identification accuracy, and genome annotation are not only determined by the chosen method but also by the specific features of the genome and repeatome being analysed.

### Transposable element library composition

We chose to analyse *D. melanogaster* alongside *Ae. albopictus* as a proof of concept, given the reliability of the extensive manual annotation available for *D. melanogaster*. Although both are classified under the term “MCTE,” the *D. melanogaster* database differs from that of *Ae. albopictus*, as the latter is not fully complete, unlike the comprehensive database described by Rech et al. (2022). This comparison allows us to assess the strengths and limitations of the ATTE method.

In *D. melanogaster*, the differences between MCTE and ATTE in TE identification are relatively minor, likely due to its smaller genome, reduced TE diversity, and lower TE content as compared with *Ae. albopictus*. Nevertheless, the same discrepancies between the two methods persist, highlighting inherent differences in their outcomes.

About 3 times more consensi was present in the ATTE library from *D. melanogaster* than in its MCTE counterpart. Furthermore, many LTR retrotransposons and DNA transposons were represented by short consensi in this ATTE library. LTR and DNA elements have a defined 5’ and 3’ ends (direct and inverted repeats, respectively) (Goubert et al. 2022; Makałowski et al. 2019; Wells and Feschotte 2020). This structure makes them relatively straightforward to detect computationally. However, identifying full-length consensus sequences remains challenging, as many insertions are fragmented or incomplete, leading automatic pipelines to often identified them as fragmented elements. Although substantial efforts by the community have been carried on in order to fix this issue as in LTR_retriver (Ou and Jiang 2018), LTR annotator (You et al., n.d.), LTRharvest (Ellinghaus, Kurtz, and Willhoeft 2008) among others, this issue is still present on modern and complex automatic pipelines such as MCHelper (S. Orozco-Arias et al. 2023). Due to their length, complex structure and propensity of their terminal repeats to recombine, the elements of this order are difficult to identify and distinguish between autonomous and non-autonomous (Flynn et al. 2020; Ji and DeWoody 2016). For what concerns the library composition of *D. melanogaster*, in spite of ATTE sequences being more numerous, the relative proportions of TE orders are not that different from those of the MCTE library.

On the contrary for *Ae. albopictus*, there are substantial differences between ATTE and MCTE libraries. For our MCTE library for *Ae. albopictus*, we identified 135 LINE elements, constituting 27.16% of the total number of consensi in the library. With these consensi, we identified 23.0% of the genome as LINEs. This percentage exceeds previous reports for this species and closely related species, such as the 12.09% from Goubert (2015) or 15.7% in *Ae. albopictus* and 14.5% in *Aedes aegypti,* the sister species of *Ae. albopictus,* reported by Melo and Wallau (2020). However, our findings align with Melo and Wallau’s observation that LINEs are the most abundant TE class.

Furthermore, we identified a high proportion of MITE elements in the MCTE library (25.96% of consensi) in *Ae. albopictus* genome. These elements are poorly characterised in current databases, despite recent efforts to classify them in insect genomes (Han et al. 2016; Chen et al. 2014; Ye, Ji, and Liang 2016). We found MITEs to be an abundant yet uncharacterized transposable element category.

Additionally, we identified elements with MITE hallmarks, such as TIRs and TSDs ranging from 800 to 3000 bp in size but devoid of any ORF. Although very similar to MITEs, the literature only confidently classifies them as such only up to a threshold of 800 bp, following the guidelines of MITEFinderII (Hu, Zheng, and Shang 2018). Therefore, longer elements were flagged as belonging to the most probable transposable origin family, labelled as “originalDNAElement_NonAutonomous” or if determining the origin family was not possible, simply as “Uncharacterized_ClassII”.

For simplicity in comparison with MCHelper annotation, these are referred to as “ClassII_others” in the figures. We also identified eight full-length insertions of *Penelope*-like elements, which have previously been reported in Culicinae mosquitoes (de Melo and Wallau 2020). These findings serve as a positive control, confirming the thoroughness of our non-exhaustive manual curation of *Ae. albopictus*.

In contrast, for the ATTE library of *Ae. albopictus*, we identified predominantly LTRs, with 73.89% of the total consensi. This might be related to detection of fragmented members of the same family of more complete LTRs as independent elements, as discussed before. As illustrated in Figure 5, LTRs, along with DNA elements (the second most abundant class for the ATTE library) and MITEs, are particularly abundant in shorter sequence lengths.

Besides these differences, we identify that due to the nature of manual curation, MCTE databases have a more detailed annotation than ATTE. For simplification of the comparison on the Figures the classification of MCTE sequences has been simplified to the superfamily. Nonetheless, the classification to family levels is more abundant in MCTE databases than ATTE ones (Supplementary Table 2).

For the *Ae. albopictus* comparison, significant differences in consensus size distribution were observed only for LTR and DNA elements. For the other TE categories, no notable differences were detected. Similarly, no significant differences were found in the *D. melanogaster* comparison. Therefore, if the objective is not to analyse TEs but to clean the assembly of them as is sometimes necessary to do on genome assembly or gene evolution analysis, an automatic method such as MCHelper seems to be a good alternative, especially on small and well characterised genomes and/or on genomes were the diversity of TE insertions is limited as in *D. melanogaster* or in humans (Mills et al. 2007).

Using our MCTE library, we identified 40.8% of the *Ae. albopictus* genome as TEs, a proportion consistent with findings from other studies. Goubert et al. (2015) found that TEs make up approximately 33.58% of the genome. More recent studies have built upon this, indicating that TEs account for over 40% of the genome (de Melo and Wallau 2020). Nonetheless, using the ATTE library we identify 73.89% of the genome as TEs. One explanation for the difference in genome coverage for the libraries of *Ae. albopictus* might be due to either the lack of exhaustiveness of the MCTE library or the use of the fully automated mode of MCHelper.

Accordingly, consensi classified as “incomplete” were kept in the final library (the consensus name has suffix “_inc”): these include all sequences that do show homology with a known TE family, but did not conserve all the expected structural or coding domain. Moreover, several “unconfirmed” sequences are also present (the consensus name has suffix “_unconfirmed”), meaning that they could only be assigned TIRs or LTRs but failed any other validation by homology or coding domain (Simon Orozco-Arias et al. 2023).

Since the *Ae. albopictus* ATTE library contains many of such sequences (5,485 incomplete and 436 unconfirmed), it is possible for the gap between our annotation and the previous ones to be caused by the detection of many motifs belonging to non-autonomous and long-extinct elements. Another possibility is for the automatic pipeline employed here may overestimate the repeat content due to an overabundance of consensi in the ATTE library. However, this is unlikely to be the case as the estimations of TE content are very consistent between the method used here and another published annotation pipeline (Baril, Imrie, and Hayward 2022) across a wide range of genome sizes (Marino et al. 2023).

On the other hand, the significant differences in the annotation outcomes between the MCTE and ATTE libraries in the two species reflect both the varying degrees of completion of the MCTE libraries and the distinct genomic architectures of the two dipterans. The *D. melanogaster* genome, at 0.18 Gb, is relatively poor in repetitive sequences, and its TE repertoire has been well-annotated, making the MCTE library a nearly exhaustive reference. In contrast, while the manual curation for *Ae. albopictus* involved three times the number of *D. melanogaster* MCTE consensi, the *Ae. albopictus* genome is 1.190 –1.275 Gb, (Palatini et al. 2020), and it is not only rich in TEs, but most of them are present in low copy numbers (Supplementary Table S3). Achieving a comprehensive description of TE diversity in the tiger mosquito through manual curation alone is an extremely labour-intensive process. However, relying solely on manual curation would result in overlooking a significant portion of the TEs present in the genome.

### How similar are the automatically generated and manually curated libraries?

For two sets of TEs libraries to be representing the TE evolutionary history in the same organism, the consensi should be highly similar sequences and represent the same variability on TE categories. To explore this idea, we decided to cluster the sequences using cd-hit-est-2d and later to compare as reciprocal species-pair -MC<->AT- using the approach presented in Flynn (2020).

Despite the varying degrees of excess sequences associated with different genomic TE densities, automatic libraries consistently contain far more “Absent” consensi compared to their manual counterparts. As previously mentioned, this may be redundant information due to the limited accuracy of the automatic “blast-extract-extend” process (Baril, Imrie, and Hayward 2022; Simon Orozco-Arias et al. 2023). However, even if the ATTE libraries retain fragmented sequences and the corresponding full-length consensi are present in the manual library, the similarity should still allow for their classification as “Fragmented” sequences. Some of the unique ATTE sequences may represent genuine “fossil” TEs that were missed -or impossible to recover- during manual curation.

Nevertheless, it is important to consider that various methods exist for curating consensi, which can result in discrepancies between libraries due to differences in multi-sequence alignment, consensus generation, sequence trimming, and other factors. This issue likely extends beyond just the automatic vs. manual comparison and applies to any comparison between different curation protocols.

Moreover, different expertises and sensitivities can also alter the outcome and lead to divergent libraries. While manual curation allows to confidently verify the validity of sequences, one disadvantage is indeed the susceptibility of such a process to human bias. In this sense, a benefit of an automatic tool is the reproducibility of the curation process.

Furthermore, the use of a single consensus sequence may be limiting, as it may fail to capture the internal variation within an entire TE family. In the future, adopting the practice of storing family information as Hidden Markov Model (HMM) profiles rather than in FASTA format could help address this limitation. HMM profiles retain information about sequence variability at each position, preserving the diversity within the TE family (J. Storer et al. 2021). Both of the libraries produced in this work -*Ae. Albopictus* MCTE and ATTE- are available on DFAM (DFAM accession number) as a multifasta files and HMM profiles based on AalbF5.

### Which method to choose then?

In most cases, TE annotation for newly sequenced genomes is delegated to either a similarity-based search (Platt, Blanco-Berdugo, and Ray 2016) or a *de novo* approach without further refinement. While this is often sufficient for basic genome masking, additional curation of TE libraries is necessary for in-depth analysis of the mobilome in the growing number of newly sequenced species. Although some examples of manual curation exist (Martelossi et al. 2023; Osmanski et al. 2023; Rech et al. 2022), carrying it out for a wide range of species or for large and TE-rich genomes like *Ae. albopictus* is time-consuming and labour-intensive. Moreover, manual curation expertise tends to be confined within certain research groups, may lack reproducibility, and, importantly, there is no single universally optimal protocol. The choice of curation method depends on the type of downstream analysis intended (Goubert et al. 2022).

In our case, the MCTE library for *Ae. albopictus* was developed with a focus on the most abundant elements showing signs of recent transposition. Manual curation was initiated to identify TE insertions potentially involved in local adaptation across different mosquito populations. Since TE polymorphisms are more likely to be associated with recent adaptations, prioritising potentially active, full-length elements was particularly relevant for this population genomics study. For such studies relying on a single reference genome, manual curation remains the best option. By contrast, manual curation would be impractical for large-scale comparative genomic analyses, such as gene evolution studies or genome assemblies, due to the time constraints and the sheer number of species typically involved.

Nonetheless, we have demonstrated that both approaches are complementary, with minimal overlap in consensi identity and contrasting strengths and weaknesses. Therefore, we provide both libraries, allowing users to decide which one (or both) they should use, depending on their specific needs.

As demonstrated in *D. melanogaster*, the TE content and composition identified using manual versus automated approaches are comparable, provided the MCTE library is comprehensive. This indicates that an automated pipeline can be appropriate for assessing TE content in a comparative framework. However, finer analysis of TE dynamics will be limited, and the results should be interpreted with caution.

## Acknowledgement

We thank Tomaso Barberis, for his work identifying consensi on *Ae. albopictus*. This work was performed using the computing facilities of the CC LBBE/PRABI.

## Data Availability

The genomic libraries generated during this study are publicly available via Zenodo. The manually curated library can be accessed at https://doi.org/10.5281/zenodo.16311754, and the automatically curated version is available at https://doi.org/10.5281/zenodo.15212853.

## Supplementary files

**Supplementary Figure 1.**
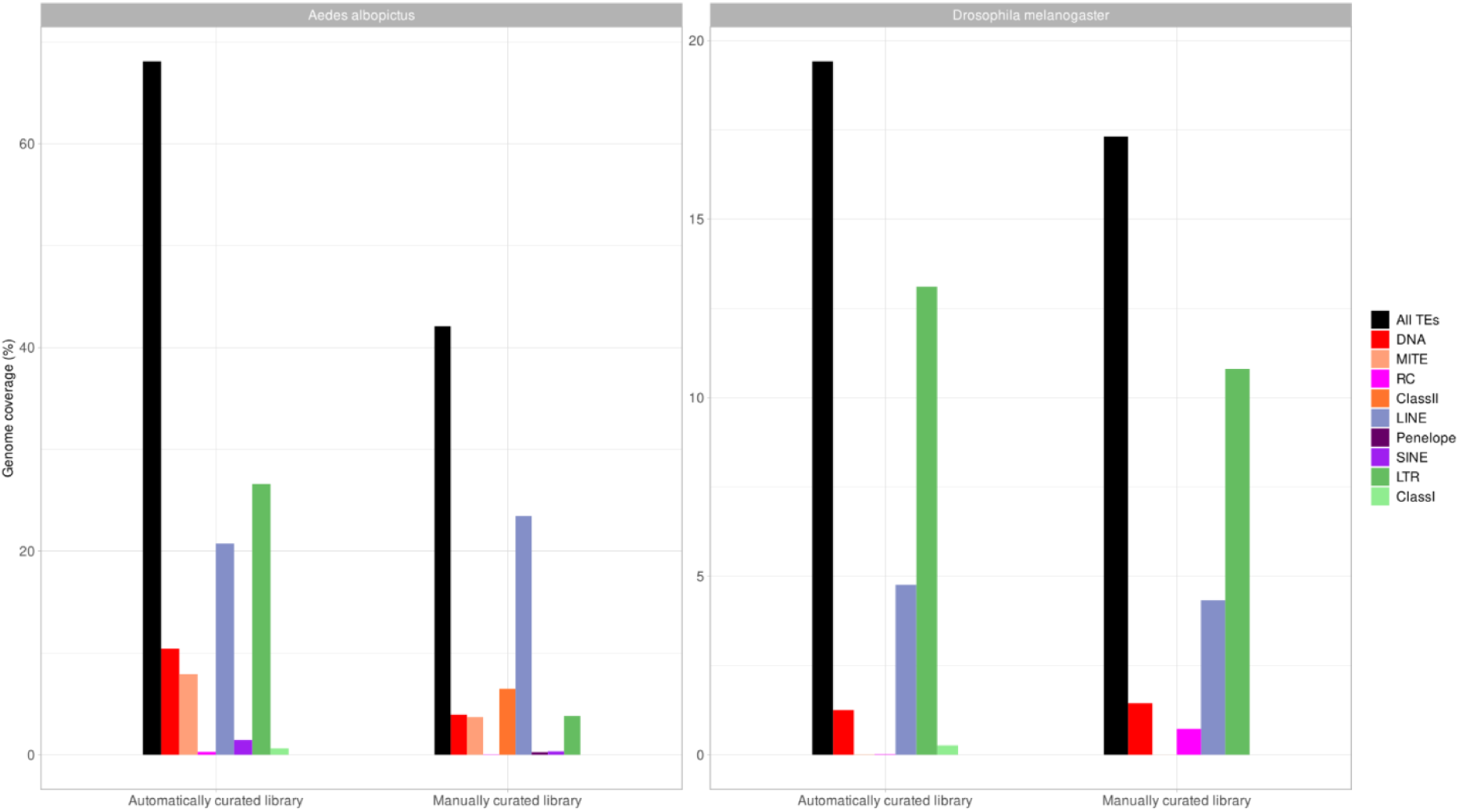
Overall (in black) and by-order genomic TE coverage identified using the automated and manual curation in *Ae. albopictus* and *D. melanogaster*. To be filled in the final figure.

### Supplementary tables

Please see:

[Supplementary Tables](https://docs.google.com/spreadsheets/d/1ctxUudPXA8xTsUX4ieqPfxiANe_7n_kVM6p1gYWx980/edit?gid=1056263486#gid=1056263486)

